# Estradiol enhances sensory-evoked dilation of cortical penetrating arterioles in a mouse model of menopause despite pre-existing metabolic dysfunction

**DOI:** 10.1101/2025.05.21.655189

**Authors:** Zachary MT Plumley, Jennifer Calvo Iglesias, Irene Fernandez Ugidos, Alice Breaux Walker, Isabela Pires dos Santos, Henry Taylor, Janet B Ruscher, Ricardo Mostany

## Abstract

Estrogen loss during menopause is a risk factor for cardiometabolic and neurovascular dysfunction, cognitive decline, and Alzheimer’s disease. Rodent studies of menopause have shown neuroprotective effects of 17β-estradiol (E2) treatment when administered within a critical period, though these benefits have not been fully demonstrated in clinical trials of hormone therapy. A proposed explanation for this mismatch is the “healthy cell bias,” where estrogen is only beneficial when initiated in physiologically intact systems. Our study investigates whether pre-existing metabolic dysfunction attenuates the effects of E2 on neurovascular coupling (NVC) in a rodent model of menopause. Female mice were fed 11 weeks of high-fat diet (HFD) or control diet (CD) to induce metabolic disease, followed by ovariectomy (OVX) and E2 or vehicle (Veh) treatment. NVC was assessed in awake mice using two-photon laser scanning microscopy of penetrating arterioles (PAs) in the somatosensory cortex, barrel field. Mice developed glucose intolerance and increased adiposity yet displayed intact NVC following HFD exposure, and subsequent OVX impaired NVC irrespective of metabolic health status. However, E2 treatment enhanced NVC responses regardless of diet relative to controls. These results indicate that estrogen’s effects on NVC are preserved in HFD+OVX conditions, which has implications for the use of hormone therapy in individuals that arrive at the menopause transition with pre-existing cardiometabolic disease.

## Introduction

Accumulating pre-clinical evidence suggests that the loss of ovarian estrogen production during the menopause transition contributes to increased risk of cardiometabolic dysfunction, age-related cognitive decline, and neurodegenerative disorders such as Alzheimer’s disease^1–3^. Indeed, estrogen is a powerful neuromodulator known to exert its protective effects at the level of neurons, the cerebral and peripheral cardiovascular system, and at the interface of these compartments: the neurovascular unit^4–7^. The critical period hypothesis of postmenopausal hormone therapy, which posits that estrogen treatment initiated within a short period following onset of menopause confers long-lasting neuroprotection, has received support in animal models of menopause^8,9^. Clinical trials of estrogen therapy, however, have yielded mixed results on cognition, ranging from null to harmful effects^10,11^. Results from the Women’s Health Initiative Memory Study (WHIMS) indicated that estrogen therapy increased risk for dementia, though treatment was initiated long after the critical period ended^12^. In the recently completed KEEPS Continuation Study, which sought to determine whether long-term benefits exist for estrogen therapy administered within the critical period, no effects of treatment were seen on cognitive performance^13^. These contrasting results have prompted new investigation into factors that may contribute to individual responsiveness to estrogen therapy.

One plausible explanation for the discrepancy between rodent and human trials of postmenopausal hormone therapy is a healthy cell bias to the beneficial effects of estrogen treatment^14–16^. Individuals may arrive at the start of the menopause transition with pre-existing cardiometabolic disorders such as obesity or diabetes, and the presence of these diseases prior to hormone therapy may limit its magnitude of effect. Such comorbidities are largely absent from the young, metabolically healthy rodents in which the neuroprotective effects of estrogen have most often been demonstrated^17^. This is important to note as estrogen itself is a regulator of energy balance and glucose homeostasis, where loss of ovarian hormones promotes adiposity and impairs insulin sensitivity, and estradiol rescues both^3,17^. Any test of the healthy cell bias must therefore contend with the possibility that estrogen’s actions on peripheral metabolism and within the brain are differentially sensitive to baseline health status. As the field moves towards precision-based medicine, it is paramount to determine whether certain postmenopausal treatments should be prioritized over others depending on a woman’s health status at menopause onset.

Neurovascular coupling (NVC), the ability of cerebral blood vessels to comply with increased neuronal energy demands, is altered in rodent models of metabolic disease^18,19^ and menopause^20^, which may have consequences for neuronal activity and downstream cognitive function. Interestingly, estrogen treatment has been shown to enhance cerebral vascular dilation^21,22^, likely through increased production of the potent vasodilator nitric oxide (NO)^23,24^. Because the neurovascular unit is tightly regulated to ensure adequate oxygen and glucose supply to the brain, probing the adaptive function of NVC during menopause may be an effective strategy to understand how the effects of postmenopausal estrogen therapy diverge in health and disease. That said, which compartment of the vascular tree should we focus on to be most sensitive to perturbations of neurovascular function? Penetrating arterioles (PAs) descend from the pial surface into the cortical parenchyma and serve as critical intermediaries between upstream pial arteries and the dense capillary network, buffering pressure gradients and distributing perfusion across cortical layers^25,26^. Additionally, PAs are the principal site of vascular resistance within the cortex and constitute a bottleneck for activity-dependent increases in blood flow^27^. Sensory-evoked dilation of individual PAs is therefore a direct and physiologically meaningful readout of changes to neurovascular function. Therefore, the current study sought to determine whether differing baseline health status prior to ovariectomy (OVX), a rodent model of menopause, affects the ability of estrogen treatment to modulate the NVC response of cortical PAs, and hypothesized that pre-existing HFD would occlude the enhancement of PA vasodilation elicited by estrogen treatment.

## Results

### Tissue weights

Uterine horn weight, body weight, heart weight, and averaged left and right kidney weights normalized to tibia length were collected at the end of each experiment to evaluate the effects of HFD, OVX, and subsequent hormone treatment on the body and peripheral tissues. Uterine horn weights of mice treated with Veh were significantly lower compared to those treated with E2 regardless of diet (Figure 1E; F(1, 75)=399.20, *p*<0.0001, 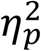=.84), indicating that OVX was sufficient to deplete endogenous estrogen and that our E2 dosage increased circulating levels of E2.

**Figure 1.**
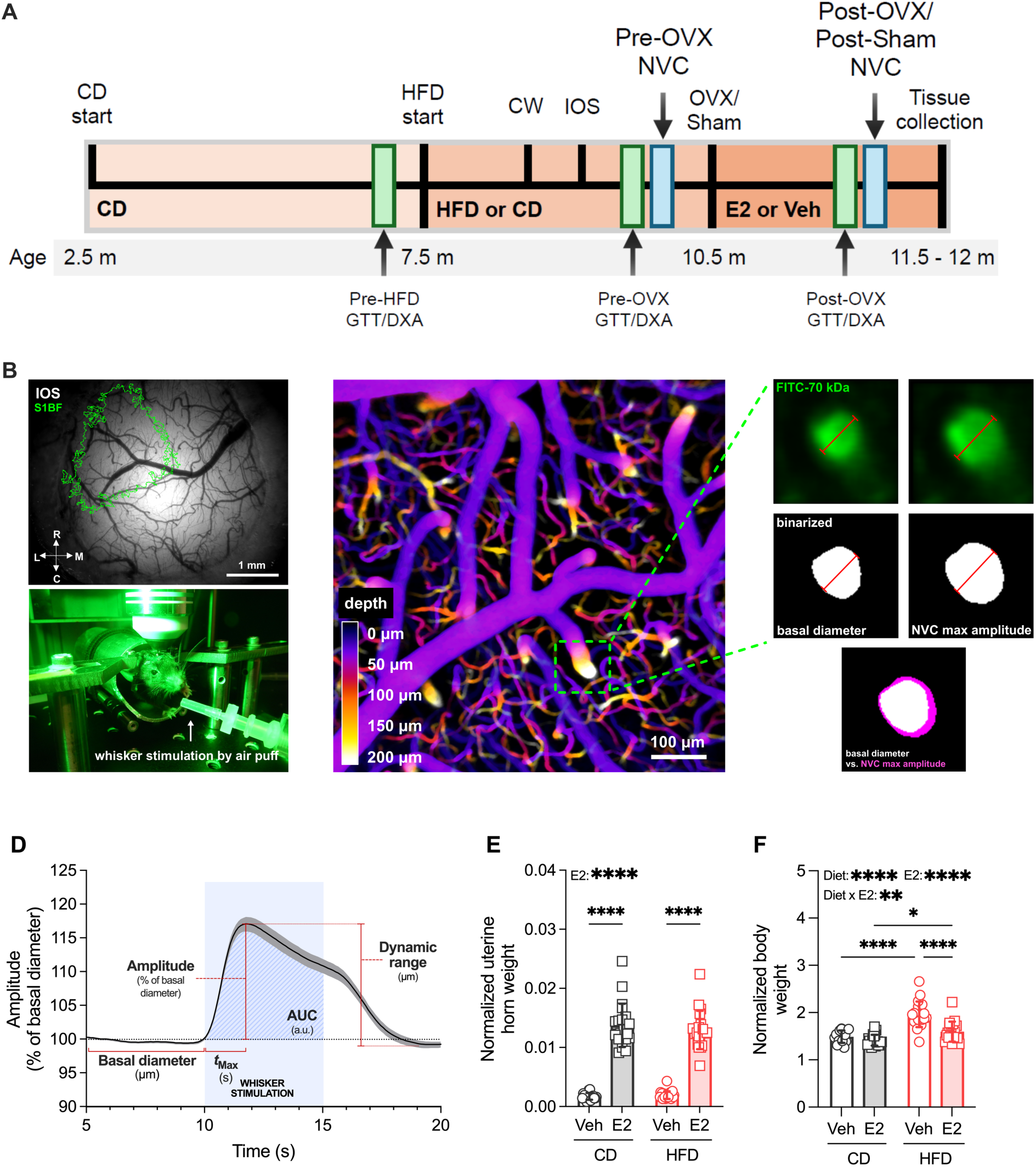
Experimental design. (A) Timeline of the study, beginning with CD start until tissue collection. (B) Bottom: microphotography of the superficial vasculature; green: highlighted location of the most caudal and medial portion of S1BF obtained by IOS imaging. Top: head-fixed awake 2PLSM set-up, with an air puff nozzle controlled by a picospritzer directed at whiskers contralateral to the cranial window. (C) Representative figure showing a pseudo-color depth-coded z-stack of the S1BF vasculature. Expanded images represent the cross-section of a PA at 50 μm depth (top row: raw images; middle row: binarized images) right before the onset of whisker stimulation (left) and at the peak of whisker stimulation-induced vasodilation (right). The bottom image shows the overlay of the binarized cross sections of the PA in basal conditions (white) and at max amplitude (magenta). (D) Example of averaged traces of NVC recordings across several mice, annotated with the NVC parameters we quantified from these traces: basal diameter (averaged PA diameter in μm from seconds 3 to 10 before whisker stimulation onset), amplitude (maximum % change in PA diameter during whisker stimulation relative to basal diameter), tMax (time in seconds after whisker stimulation onset to reach maximum amplitude), AUC (area under the curve calculated during whisker stimulation, blue-striped area), dynamic range (maximum absolute change in diameter in μm across the time series), and basal diameter (μm). Blue-shaded area indicates the duration of whisker stimulation. (E) Comparison of uterine horn weights (g/tibia length in μm) to 1) verify ovarian hormone depletion following OVX and 2) assess bioavailability of E2 silastic capsule treatment (n=16 CD/Veh-treated mice, n=21 CD/E2-treated mice, n=22 HFD/Veh-treated mice, and n=20 HFD/E2-treated mice). (F) Comparison of terminal body weight (g/tibia length in μm) (n=16 CD/Veh-treated mice, n=20 CD/E2-treated mice, n=21 HFD/Veh-treated mice, and n=20 HFD/E2-treated mice). *p<0.05; **p<0.01; ****p<0.0001. Veh: vehicle; E2: estradiol; CD: control diet; HFD: high-fat diet; Pre-OVX/Sham: pre-ovariectomy/sham timepoint; Post-OVX/Sham: post-ovariectomy/sham timepoint; CW: chronic cranial window implantation; IOS: intrinsic optical signal imaging; NVC: *in vivo* awake two-photon laser scanning microscopy of neurovascular coupling of penetrating arterioles; S1BF: primary somatosensory cortex barrel field; FITC: 70 kDa fluorescent isothiocyanate conjugated dextran used for labelling serum to visualize vessel diameter change.

Mice given HFD weighed more than CD-fed mice at study end (Figure 1F; F(1, 73)=52.93, *p*<0.0001, 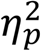=.42), and mice treated with E2 weighed less than those given Veh (F(1, 73)=24.15, *p*<0.0001, 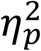=.25). Interestingly, we noted an interaction where the effect of E2 on reduction of body weight (relative to Veh) was more profound in HFD-vs. CD-fed mice (F(1, 73)=10.62, *p*=0.0017, 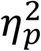=.13). HFD-fed mice showed increased heart weight (i.e. cardiac hypertrophy) relative to those fed CD regardless of hormone treatment (Figure S1A; F(1, 74)=8.07, *p*=0.0058, 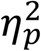=.10), while kidney weights were not different between any of the groups (Figure S1B).

### Dual energy x-ray absorptiometry (DEXA) scans

Body fat and lean mass of each mouse were measured by DEXA scans performed at the Pre-HFD, Pre-OVX, and Post-OVX timepoints. At study start, no differences were found in fat mass between mice that received CD throughout the study or those assigned to later receive HFD (Figure 2A). At the Pre-OVX timepoint, mice fed HFD for 11 weeks showed increased adiposity compared to CD-fed mice (Figure 2B; CD *vs*. HFD; 5.98 ± 2.07 *vs.* 16.30 ± 4.11; *p*<0.0001, *d*=0.15). Following ovariectomy and short-term E2 or Veh treatment, we observed that HFD-fed mice had elevated body fat mass compared to those given CD (Figure 2C; F(1, 91)=159.30, *p*<0.0001, 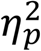=.64), mice treated with E2 had lower body fat mass relative to Veh (Figure 2C; F(1, 91)=46.46, *p*<0.0001, 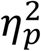=.34), and identified a significant interaction between diet and E2, where HFD-fed mice given E2 had less body fat compared to the Veh group, while there were no significant effects of E2 in the CD condition (Figure 2C-D; F(1, 91)=14.49, *p*=0.0003, 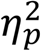=.14). Importantly, no effects of HFD or E2 on lean mass were observed at the Pre-HFD, Pre-OVX, or Post-OVX timepoints (Figure 2E-H), which suggests that group differences seen in body weight at end of study were mediated primarily by changes in fat mass driven by diet and E2 treatment.

**Figure 2.**
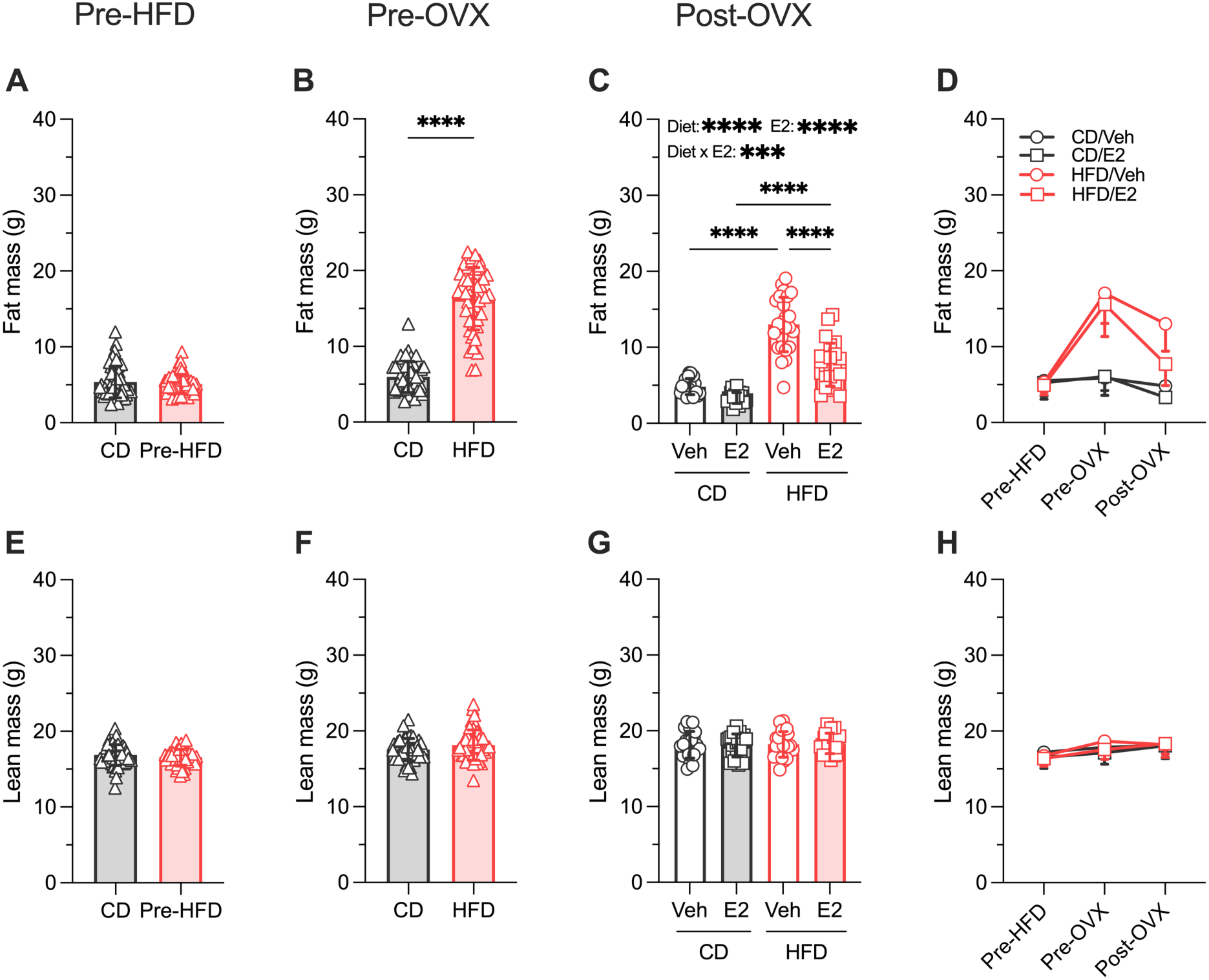
Body composition assessment by dual x-ray absorptiometry (DEXA) scans. (A-D) Fat mass and (E-H) lean mass computed from DEXA scan analysis at Pre-HFD (A,E; n=51 CD-fed mice vs. 55 HFD-fed mice), Pre-OVX (B,F; n=45 CD-fed mice vs. 46 HFD-fed mice), and Post-OVX (C,G; n=22 CD/Veh-treated mice, 23 CD/E2-treated mice, 25 HFD/Veh-treated mice, and 25 HFD/E2-treated mice). ***p<0.001; ****p<0.0001. Pre-HFD: pre-high fat diet timepoint; Pre-OVX: pre-ovariectomy timepoint; Post-OVX: post-ovariectomy timepoint; Veh: vehicle; E2: estradiol; CD: control diet; HFD: high-fat diet.

### Glucose tolerance test (GTT)

In addition to the DEXA scans, we analyzed fasting blood glucose levels and calculated area under the curve (AUC) from glucose tolerance tests performed at the Pre-HFD, Pre-OVX, and Post-OVX timepoints to determine the effects of HFD on glucose metabolism, and whether pre-existing HFD-induced metabolic disease moderates the effects of E2 treatment on glucose homeostasis following ovariectomy. As observed with the DEXA data, no differences were found in basal glucose levels or AUC between mice assigned to later receive HFD or those continuing CD for the remainder of the study (Figures 3A-C). At the Pre-OVX timepoint, HFD-fed mice showed elevated basal glucose levels (Figure 3D; CD *vs*. HFD; 170.60±26.59 *vs.* 196.50±35.63; *p*=0.0004, *d*=0.82) and AUC measurements (Figures 3E-F; CD *vs*. HFD; 27727.00±11605.00 *vs.* 43095.00±4331.00; *p*<0.0001, *d*=1.75) compared to CD-fed mice, suggesting that exposure to HFD impaired systemic glucose regulation. At the Post-OVX timepoint, mice treated with E2 had lower basal glucose levels compared to their Veh-treated counterparts (Figure 3G; F(1, 88)=50.68, *p*<0.0001, 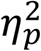=.37). We again identified a diet x E2 treatment interaction where HFD-fed mice given E2 yielded significantly lower basal glucose levels than their HFD-Veh counterparts, while the effects of E2 were less pronounced in the CD-fed mice (Figure 3G; F(1, 88)=16.02, *p*<0.0001, 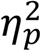=.15). Our AUC measurements showed a similar pattern: following the glucose challenge, E2-treated mice had lower AUC compared to Veh (Figure 3H-I; F(1, 88)=21.12, *p*<0.0001, 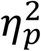=.19), HFD mice had higher AUC compared to CD-fed mice (Figure 3H-I; F(1, 88)=64.84, *p*<0.0001, 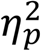=.42), and a significant interaction revealed the effect of E2 treatment on AUC was more pronounced in HFD-fed mice than in those fed CD, relative to Veh (Figure 3H-I; F(1, 88)=6.25, *p*<0.0143, 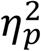=0.07). Taken together with the body weight and DEXA scan measurements, our data indicate that 11 weeks of exposure to a HFD (with 59% kcal from fat) reliably induces a systemic metabolic disease in middle-aged female mice, and that short-term E2 treatment following subsequent mid-life OVX surgery is sufficient to rescue this phenotype.

**Figure 3.**
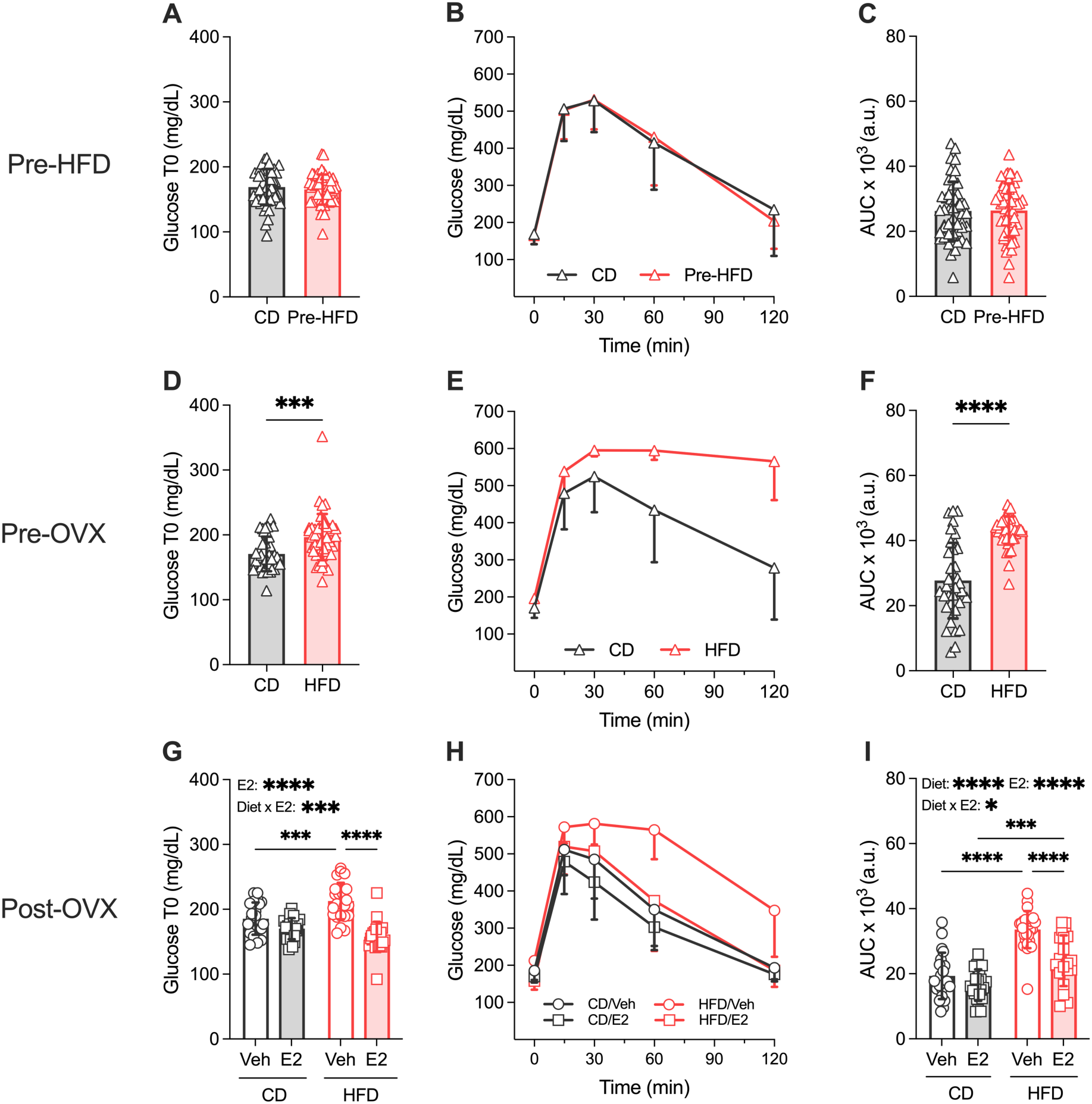
Glucose tolerance tests. (A,D,G) Fasting glucose levels, (B,E,H) blood glucose curves, and (C,F,I) AUC computed from GTT at Pre-HFD (A-C; n=47 CD-fed mice vs. n=52 HFD-fed mice), Pre-OVX (D-F; n=38 CD-fed mice vs. n=47 HFD-fed mice), and Post-OVX (G-I; n=22 CD/Veh-treated mice, n=22 CD/E2- treated mice, n=25 HFD/Veh-treated mice, n=23 HFD/E2-treated mice). *p<0.05; ***p<0.001; ****p<0.0001. Pre-HFD: pre-high fat diet timepoint; Pre-OVX: pre-ovariectomy timepoint; Post-OVX: post-ovariectomy timepoint; Veh: vehicle; E2: estradiol; CD: control diet; HFD: high-fat diet; AUC: area under the curve; Glucose T0: fasting glucose levels measured before glucose injection.

### Neurovascular coupling: Pre-OVX CD vs. HFD

We next sought to evaluate the effects of the HFD-induced metabolic disease on awake NVC responses to air puff whisker stimulation prior to OVX (Figure 4A). Imaged PAs ranged between 28-63 µm in basal diameter (Figure 4B), which were not significantly different between CD and HFD-fed mice (Figure 4C). We did not observe any differences in amplitude, t_Max_, AUC, or dynamic range of analyzed PAs from CD and HFD-fed mice (Figure 4D-G). Additionally, we found no correlations between a given PA’s basal diameter to its respective amplitude, t_Max_, AUC, or dynamic range in mice receiving HFD or CD (Figure 4H-K). These findings indicate that in the intact middle-aged female mouse, cortical PAs resist perturbation from systemic HFD-induced metabolic dysfunction, and that the amplitude of the NVC response is not dictated by basal diameter.

**Figure 4.**
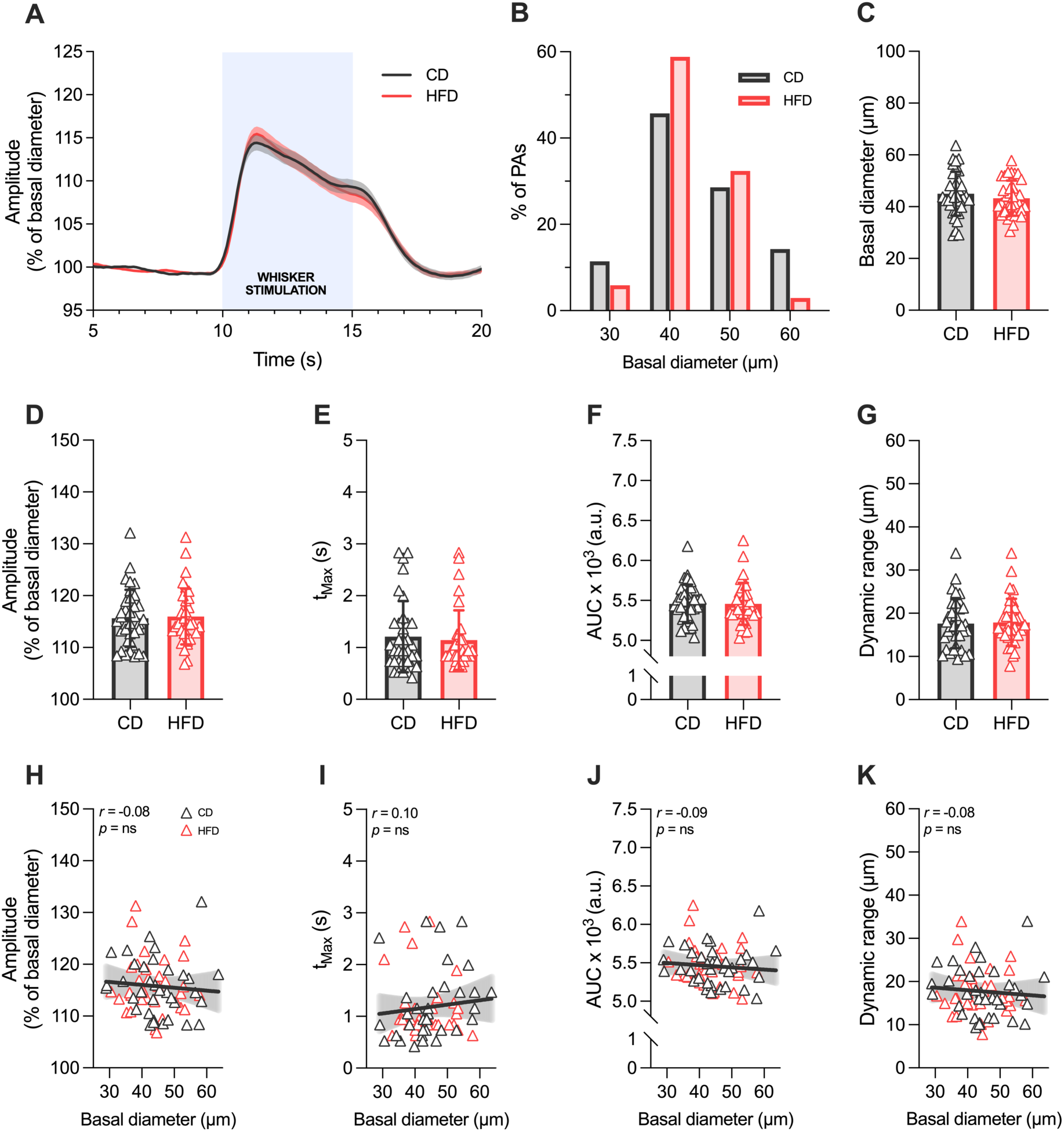
Neurovascular coupling: Pre-OVX CD vs. HFD. (A) Averaged traces of whisker stimulation-evoked PA vasodilation in CD-fed mice (n=35 PAs from N=18 mice) vs. HFD-fed mice (n=34 PAs from N=18). (B) Distribution of basal diameter values of all PAs classified into 10 μm diameter bins. (C-G) Comparison of (C) basal diameter, (D) amplitude, (E) tMax, (F) AUC, and (G) dynamic range in CD-fed mice (n=35 PAs from N=18 mice) vs. HFD-fed mice (n=34 PAs from N=18) prior to OVX. (H-K) Correlation plots of a given PA’s (H) amplitude, (I) tMax, (J) AUC, and (K) dynamic range to their respective basal diameter in 18 CD-fed mice (n=35 PAs from N=18 mice) vs. HFD-fed mice (n=34 PAs from N=18) prior to OVX. CD: control diet; HFD: high-fat diet.

### Neurovascular coupling: Post-Sham vs. Post-OVX

Next we wanted to determine whether OVX in middle age is sufficient to impair PA NVC, and further, whether the double hit of HFD+OVX would result in a worsened phenotype. Measurements of PA basal diameter and NVC responses were performed 4-6 weeks following either OVX+Veh capsule implantation or Sham procedures (Figure 1A, 5A). Imaged PAs again ranged between 28-63 µm in basal diameter (Figure 5B), and there was no effect of diet or OVX on basal diameter (Figure 5C). We observed that OVX decreased the amplitude of the NVC response (Figure 5A, 5D; F_(1, 85)_ = 4.40, *p*=0.0390, 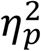=0.05), did not affect t_Max_ or AUC (Figure 5E-F), and restricted dynamic range (Figure 5G; F_(1, 85)_ = 4.19, *p*=0.0438, 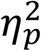=0.05). In agreement with the Pre-OVX results, we did not observe an effect of HFD on any of our NVC parameters (Figure 5D-G).

**Figure 5.**
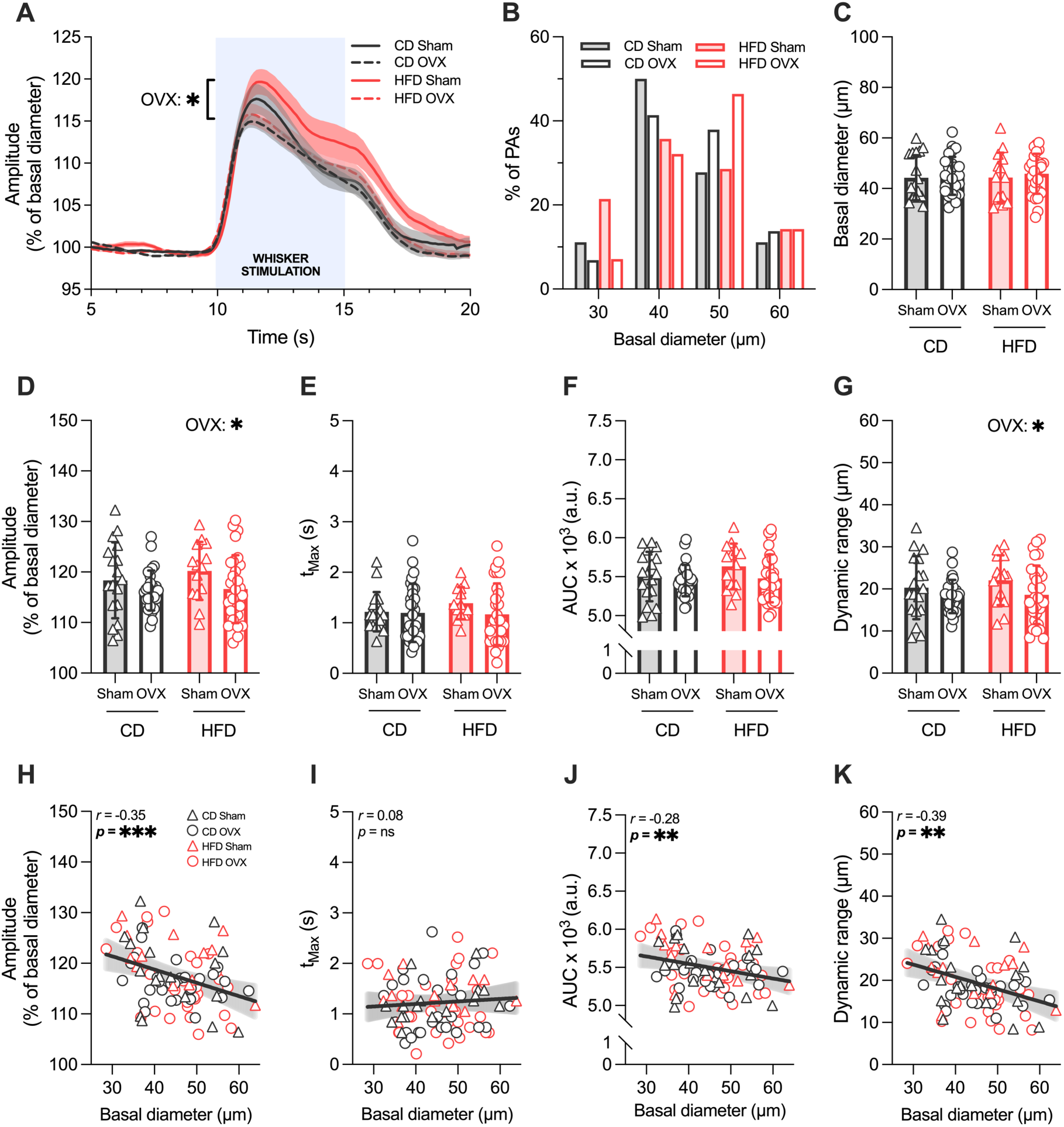
Neurovascular coupling: Sham vs. OVX. (A) Averaged traces of whisker stimulation-evoked PA vasodilation in CD/Sham mice (n=18 PAs from N=9 mice), CD/OVX Veh-treated mice (n=28 PAs from N=15 mice), HFD/Sham mice (n=14 PAs from N=7 mice), and HFD/OVX Veh-treated mice (n=29 PAs from N=15 mice). (B) Distribution of basal diameter values of all PAs classified into 10 μm diameter bins. (C-G) Comparison of (C) basal diameter, (D) amplitude, (E) tMax, (F) AUC, and (G) dynamic range of PAs from CD/Sham mice (n=18 PAs from N=9 mice), CD/OVX Veh-treated mice (n=28 PAs from N=15 mice), HFD/Sham mice (n=14 PAs from N=7 mice), and HFD/OVX Veh-treated mice (n=29 PAs from N=15 mice). (H-K) Correlation plots of a given PA’s (H) amplitude, (I) tMax, (J) AUC, and (K) dynamic range to their respective basal diameter in CD/Sham mice (n=18 PAs from N=9 mice), CD/OVX Veh-treated mice (n=28 PAs from N=15 mice), HFD/Sham mice (n=14 PAs from N=7 mice), and HFD/OVX Veh-treated mice (n=29 PAs from N=15 mice). *p<0.05; **p<0.01; ***p<0.001. CD: control diet; HFD: high-fat diet.

Although basal diameter was not perturbed by either HFD or OVX, we found it was negatively correlated with amplitude (Figure 5H; r=-0.35, n=89, p=0.0007), not correlated with tMax (Figure 5I), and negatively correlated with both AUC (Figure 5J; r=-0.28, n=89, p=0.0074) and dynamic range (Figure 5K; r=-0.39, n=89, p=0.0002) across treatment groups, indicating that OVX may preferentially impair larger PAs and/or recruit larger responses from smaller PAs. Taken together, these results indicate that the loss of ovarian hormones from OVX impairs the magnitude of awake sensory-evoked vasodilation, and that OVX may alter the contributions of differently-sized PAs to amplitude change.

### Neurovascular coupling: Post-OVX Veh vs. E2

Since we determined OVX is sufficient to impair the amplitude and dynamic range of PA dilation, we sought to evaluate whether this blunted NVC response was specifically due to the loss of endogenous E2 (an ovarian hormone known to prime experience-dependent vasodilation) and whether E2 treatment can rescue NVC following OVX. Importantly, despite the lack of a direct HFD effect on PA basal diameter and NVC parameters in prior experiments, we wanted to determine whether previous HFD would limit the extent of E2’s dilation-potentiating effects in PAs. NVC measurements were performed 4-6 weeks following OVX and either E2 or Veh capsule implantation (Figure 1A). Imaged PAs ranged between 28-62 µm in basal diameter (Figure 6B), and again there was no effect of diet or E2 on basal diameter (Figure 6C). Our data shows that E2 significantly increased PA amplitude (Figure 6A, 6D; F_(1, 108)_=14.09, *p*=0.0003, 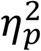=.12), did not affect t_Max_ (Figure 6E), and increased both AUC (Figure 6F; F_(1, 108)_=10.76, *p*=0.0014, 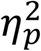=.09) and dynamic range (Figure 6G; F_(1, 108)_=13.22, *p*=0.0004, 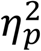=.11). Though no interaction effect between HFD+E2 was observed, post hoc analyses revealed that the main effect of E2 was primarily driven through CD-fed mice in amplitude (Figure 6D; CD+Veh *vs*. CD+E2; 116.30±3.97 *vs.* 121.70±6.66; *p*=0.0020, *d*=0.33), AUC (Figure 6F; CD+Veh *vs*. CD+E2; 5.45±0.20 *vs.* 5.68± 0.26; *p*=0.0048, *d*=0.63), and dynamic range (Figure 6G; CD+Veh *vs*. CD+E2; 18.22± 3.95 *vs.* 23.64± 6.93; *p*= 0.0027, *d*=0.67). Interestingly, HFD prior to OVX shortened t_Max_, though this effect was relatively small (Figure 6E; F_(1, 108)_=5.00, *p*=0.0275 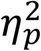=.04) and yielded no effects on amplitude, AUC, or dynamic range. This effect of HFD was likely driven by prolonged t_Max_ in the CD+E2 group relative to HFD+E2 (Figure 6E; CD+E2 *vs*. HFD+E2; 1.54±0.48 *vs.* 1.13±0.37; *p*=0.0159, *d*=0.57), consistent with a more pronounced effect of E2 on the amplitude, AUC, and dynamic range in CD vs. HFD-fed mice.

**Figure 6.**
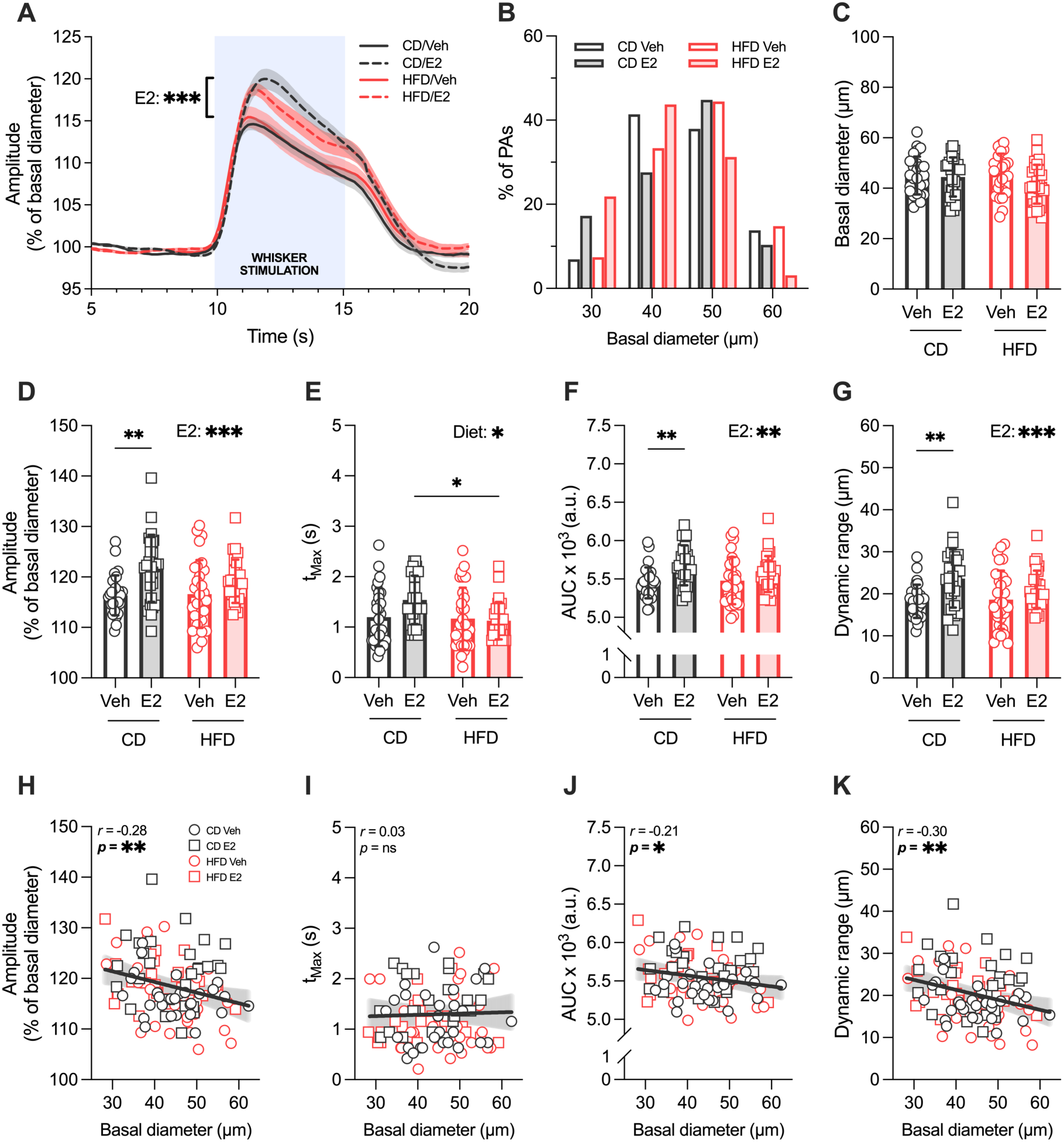
Neurovascular coupling: Post-OVX Veh vs. E2. (A) Averaged traces of whisker stimulation-evoked PA vasodilation in CD/OVX Veh-treated mice (n=28 PAs from N=15 mice), CD/OVX E2-treated mice (n=27 PAs from N=14 mice), HFD/OVX Veh-treated mice (n=29 PAs from N=15 mice), and HFD/OVX E2-treated mice (n=28 PAs from N=16 mice). (B) Distribution of basal diameter values of all PAs classified into 10 μm diameter bins. (C-G) Comparison of (C) basal diameter, (D) amplitude, (E) tMax, (F) AUC, and (G) dynamic range of PAs from CD/OVX Veh-treated mice (n=28 PAs from N=15 mice), CD/OVX E2-treated mice (n=27 PAs from N=14 mice), HFD/OVX Veh-treated mice (n=29 PAs from N=15 mice), and HFD/OVX E2-treated mice (n=28 PAs from N=16 mice). (H-K) Correlation plots of a given PA’s (H) amplitude, (I) tMax, (J) AUC, and (K) dynamic range to their respective basal diameter in CD/OVX Veh-treated mice (n=28 PAs from N=15 mice), CD/OVX E2-treated mice (n=27 PAs from N=14 mice), HFD/OVX Veh-treated mice (n=29 PAs from N=15 mice), and HFD/OVX E2-treated mice (n=28 PAs from N=16 mice). *p<0.05; **p<0.01; ***p<0.001. Veh: vehicle; E2: estradiol; CD: control diet; HFD: high-fat diet; AUC: area under the curve.

Looking at the relationship of PA size to NVC, we found that basal diameter was negatively correlated with amplitude (Figure 6H; r=-0.28, n=112, p=0.0030), not correlated with tMax (Figure 6I), and correlated with both AUC (Figure 6J; r=-0.21, n=112, p=0.0281) and dynamic range (Figure 6K; r=-0.30, n=112, p=0.0011) across treatment groups, indicating that E2 treatment after OVX restores stochastic basal diameter x NVC dynamics.

The potentiating effect of E2 treatment on amplitude, AUC, and dynamic range despite pre-existing metabolic disease is consistent with findings from our Post-Sham vs. Post-OVX experiments, where a decrease in production of ovarian hormones due to OVX resulted in a blunted NVC response. Since we did not observe a difference in t_Max_ between E2 and Veh-treated mice, our data suggests that E2 preferentially enhances and sustains the NVC response while exerting less of an effect on response kinetics, largely independent of the influence of baseline health status.

### Neurovascular coupling x basal diameter: Post-Sham and Post-OVX

Given our observation that basal diameter was negatively correlated with NVC amplitude, AUC, and dynamic range in our Post-Sham vs. Post-OVX and Post-OVX E2 vs. Veh experiments, but not in Pre-OVX, we wondered if it was possible that ovarian hormone status influences NVC differently depending on relative PA size? To answer this question, we analyzed the correlation between PA basal diameter and NVC amplitude in Sham-operated, OVX Veh, and OVX E2 mice. In Sham-operated mice, we found that basal diameter was not correlated with amplitude (Figure 7A). However, in OVX mice basal diameter was negatively correlated with amplitude (Figure 7A-B; *r*=-0.40, *n*=57, *p*=0.0022), indicating OVX may preferentially impair larger PAs and/or recruit larger responses from smaller PAs. This correlation was not observed in OVX E2 mice, indicating a return to apparently stochastic basal diameter/NVC response dynamics (Figure 7B).

**Figure 7.**
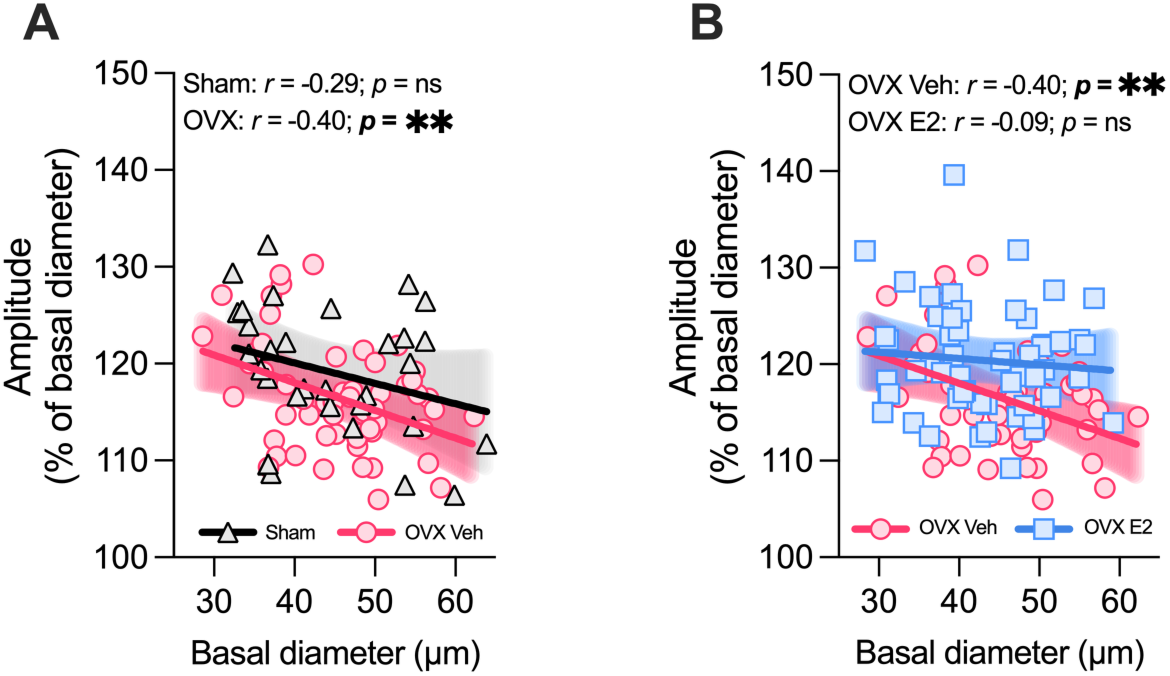
Neurovascular coupling x basal diameter: Post-Sham and Post-OVX. (A) Correlation plots of a given PA’s amplitude to their respective basal diameter in CD+HFD Sham mice (n=32 PAs from N=16 mice) and CD+HFD OVX/Veh-treated mice (n=57 PAs from N=30 mice), and (B) in CD+HFD OVX/Veh-treated mice (n=57 PAs from N=30 mice) and CD+HFD OVX/E2-treated mice (n=55 PAs from N=30 mice). **p<0.01. Veh: vehicle; E2: estradiol; CD: control diet; HFD: high-fat diet

To account for the possibility that intrinsic functional properties related to a given PA’s basal diameter might independently affect or interact with NVC responses and therefore bias our readouts, we performed a multiple linear regression analysis to evaluate the effects of HFD, OVX, and E2 treatment on maximum absolute amplitude while controlling for effects solely due to basal diameter. In the Post-Sham vs. Post-OVX experiment, HFD, OVX, nor the interaction of basal diameter and OVX were significant predictors of absolute amplitude independent of basal diameter alone (Table 1). This indicates that stochastic properties of differently sized PAs cannot be separated from the magnitude of effect HFD and OVX impart on NVC. For the Post-OVX E2 vs. Veh experiment, we found E2 and HFD were not independently significant predictors of absolute amplitude controlling for the effects of basal diameter. Interestingly however, we did find a significant interaction between basal diameter and E2 treatment, suggesting that a given PA’s size directly dictates the magnitude of enhancement E2 imparts on NVC (Table 2). Taken together, these results demonstrate that the loss of ovarian hormones contributes to dysregulated PA-NVC dynamics, where larger PAs typically elicit a truncated amplitude response, and that E2 treatment normalize stochastic PA-NVC dynamics.

**Table 1.** Multiple regression analysis: Post-Sham vs. Post-OVX predictors of absolute NVC amplitude. Multiple regression analysis indicating the predictive power of basal diameter, CD vs. HFD, Sham vs. OVX, and the interaction of basal diameter and Sham vs. OVX on absolute NVC amplitude (i.e. the maximum whisker stimulation-evoked diameter of each PA in μm). *df*: degrees of freedom; *B*: unstandardized regression coefficient; *SE*: standard error; 95% *CI*: 95% confidence interval; *LL*: lower limit; *UL*: upper limit; *R*: relative strength of correlation between observed vs. predicted values from model; *R^2^*: fraction of absolute amplitude explained by predictors combined; *R^2^_adj_*: fraction of absolute amplitude explained by predictors combined adjusted for redundancy in predictors.

| Effects | **df** | ***B*** | ***SE*** | **95% CI** |  | ***p*** |
|---|---|---|---|---|---|---|
|  |  |  |  | ***LL*** | ***UL*** |  |
| Regression Model | 4 |  |  |  |  |  |
| Intercept |  | 4.09 | 2.26 | -0.399 | 8.57 | 0.0736 |
| Basal diameter | 1 | 1.09 | 0.0496 | 0.993 | 1.19 | <0.0001 |
| HFD (0=CD; 1=HFD) | 1 | 0.360 | 0.538 | -0.710 | 1.43 | 0.5051 |
| OVX (0=Sham; 1=OVX) | 1 | 1.00 | 3.03 | -5.02 | 7.03 | 0.7409 |
| Basal diameter x OVX | 1 | -0.0469 | 0.0665 | -0.179 | 0.0853 | 0.4823 |
| Residual | 84 |  |  |  |  |  |
| Total | 88 |  |  |  |  |  |

*R*=0.962, *R*<sup>2</sup>=0.926, *R*<sup>2</sup><sub>adj</sub>=0.922, F<sub>(4, 84)</sub>=261, *p*<0.0001

**Table 2.** Multiple regression analysis: Post-OVX E2 vs. Veh predictors of absolute NVC amplitude. Multiple regression analysis indicating the predictive power of basal diameter, CD vs. HFD, Veh vs. E2, and the interaction of basal diameter and Veh vs. E2 on absolute NVC amplitude. *df*: degrees of freedom; *B*: unstandardized regression coefficient; *SE*: standard error; 95% *CI*: 95% confidence interval; *LL*: lower limit; *UL*: upper limit; *R*: relative strength of correlation between observed vs. predicted values from model; *R^2^*: fraction of absolute amplitude explained by predictors combined; *R^2^_adj_*: fraction of absolute amplitude explained by predictors combined adjusted for redundancy in predictors.

| Effects | **df** | ***B*** | ***SE*** | **95% CI** |  | ***p*** |
|---|---|---|---|---|---|---|
|  |  |  |  | ***LL*** | ***UL*** |  |
| Regression Model | 4 |  |  |  |  |  |
| Intercept |  | 5.42 | 1.86 | 1.74 | 9.10 | 0.0043 |
| Basal diameter | 1 | 1.05 | 0.0402 | 0.969 | 1.13 | <0.0001 |
| HFD (0=CD; 1=HFD) | 1 | -0.585 | 0.439 | -1.46 | 0.285 | 0.1851 |
| E2 (0=Veh; 1=E2) | 1 | -3.87 | 2.57 | -8.97 | 1.23 | 0.1352 |
| Basal diameter x E2 | 1 | 0.126 | 0.0573 | 0.0120 | 0.239 | 0.0306 |
| Residual | 107 |  |  |  |  |  |
| Total | 111 |  |  |  |  |  |

*R*=0.967, *R*<sup>2</sup>=0.935, *R*<sup>2</sup><sub>adj</sub>=0.933, F<sub>(4, 107)</sub>=386, *p*<0.0001

## Discussion

This is the first study to our knowledge to evaluate whether a healthy cell bias exists for E2 to exert regulatory control over functional hyperemia in a rodent model of menopause. Using *in vivo* 2PLSM, we quantified changes in the dynamics of cortical PAs, and found that 11 weeks of HFD does not affect NVC. Our data provide strong evidence for the role of E2 in influencing the magnitude of the NVC response at mid-life, and that this E2 action persists in the context of health and disease. We also find that OVX impairs dynamics of larger vessels, which is rescued by E2 treatment. These results suggest that the neurovascular unit retains sensitivity to estrogen signaling during female aging despite compromised peripheral glucose regulation.

Unlike capillaries, which rely on local signaling to redistribute blood flow, or pial arterioles that direct more broad hemodynamic changes^27^, PAs are critical intermediaries between upstream pial arteries and the dense capillary network, buffering pressure gradients and directing perfusion across cortical layers^25,26^. Despite this, it is plausible that PAs may exhibit less metabolic disease-induced dysfunction relative to other vascular compartments. One recent study showed that exposure to a HFD increased both endothelial nitric oxide synthase (eNOS) and astrogliosis, indicating a compensatory mechanism for increased vasodilation despite increased inflammation^38^. Another group using the Akita mouse model of type 1 diabetes found no differences in sensory-evoked vasodilation of PAs in the S1BF^39^. A prior study from our laboratory echoed these findings; in a streptozotocin (STZ) model of type 1 diabetes, no effects of chronic hyperglycemia were found on basal diameter or NVC responses of S1BF PAs^33^. Our results support this interpretation: despite the presence of glucose intolerance and increased adiposity of HFD-fed mice, basal diameter and NVC responses were not affected. It is also possible that capillaries could be the prime target of the effects of HFD+OVX given their reliance on pericyte-endothelial cell signaling cascades which are sensitive to metabolic insult^40,41^. Future work could determine if there is differential compartmental vulnerability to HFD+OVX in the vascular tree.

Despite preserved NVC with pre-existing metabolic disease, OVX impaired the NVC magnitude which is consistent with previous studies demonstrating estrogen loss alters the function of cerebral blood vessels. OVX has been shown to reduce eNOS expression, linked to concurrent changes in caveolin-1 (the endogenous inhibitor of eNOS) in the brain^42^. Since eNOS-derived NO is a mediator of NVC, its downregulation following OVX may contribute to the decrease in amplitude and dynamic range we observed. Interestingly, caveolin-1 functions not only to limit eNOS activity but also to promote NVC in the cerebral cortex. In a recent study, cortical arteriole endothelial cell caveolae were found to relay vasodilatory signals to smooth muscle cells, driving the NVC response independently from the eNOS pathway^43^.

Beyond eNOS, OVX exerts broader effects on NVC. OVX increases arterial stiffness and downregulates genes for peripheral vessel contractility, which is reversible through activation of the G-protein coupled estrogen receptor (GPER)^44^, raising the possibility that similar shifts occur in cortical PAs that contribute to elevated myogenic tone documented in these vessels^45^. Increased tone may narrow the dilatory reserve for sensory-evoked responses, consistent with the selective impairment of NVC magnitude we observed. Additionally, the 4-vinylcyclohexene diepoxide (VCD) model of menopause results in impaired large-conductance Ca^2+^ activated potassium (BK_Ca_) channel function in arterioles while blunting functional hyperemia^20^, representing an additional possible mechanism behind OVX’s impairments of NVC. That the effects of OVX on NVC did not depend on prior HFD exposure suggests ovarian hormones and metabolic disease alter NVC through non-overlapping pathways.

E2 enhanced the NVC response following OVX regardless of metabolic disease, which is consistent with previous work demonstrating that E2 taps several genomic and non-genomic pathways to increase vasodilation, resulting in increased endothelial NO release, neuron-astrocyte signaling, and potassium channel activation as discussed before^6,7,20,23^. At the receptor level, rapid effects of E2 in the cerebrovascular endothelium are primarily mediated by GPER^4^, which induces vasodilation of PAs through both eNOS and direct action on vascular smooth muscle^46^. GPER signaling may represent an additional pathway through which E2 primes PAs for enhanced NVC. Mechanistically, E2 reduces myogenic tone in the cerebral vasculature through NO-dependent vessel relaxation, wherein estrogen receptor alpha (ERα) upregulation in endothelial cells increases eNOS expression^24,47–49^. These receptor-level changes resulting in endothelial NO-mediated enhancements in vasodilation likely underlie our observed effects of E2 on NVC magnitude without significantly affecting t_Max_, which may be more dependent on upstream neuronal or astrocytic calcium signaling kinetics. Interestingly, t_Max_ was also unaffected by sustained hyperglycemia in our previous work using the STZ model^33^. The dissociation between magnitude and kinetics-related NVC parameters suggests they may be independently regulated within the neurovascular unit.

How do our findings compare to other investigations into the healthy cell bias of estrogen action in different systems of the brain? Several recent findings from our multi-laboratory collaborative research team have contributed to bridging this knowledge gap. One study demonstrated that the cognitive benefits of E2 during female aging are independent from its benefits in rescuing effects of OVX on cardiometabolic health^50^. Using the same model of metabolic disease, another study found HFD attenuated the beneficial effects of E2 on spatial memory task performance and blunted E2-mediated increases in hippocampal long-term potentiation^29^. However, a third study found increased excitability of PVN liver-related neurons in HFD+OVX mice, which was actually rescued by E2^30^. Lastly, another report found that E2 following OVX could not rescue aortic oxidative stress in angiotensin-induced hypertensive mice. Taken together, if a healthy cell bias of estrogen exists in the brain, it may be better viewed as compartment-specific, where neuronal or vascular targets of E2 may be differentially vulnerable to the double hit of HFD+OVX.

Across experimental groups, we observed an inverse relationship between PA basal diameter and magnitude of NVC response, with smaller PAs exhibiting greater NVC magnitude despite no effect of HFD or E2 on basal diameter itself. A similar correlation has been documented in PAs of the mouse S1BF across both healthy and diabetic animals^39^. We found the significant correlation between basal diameter and NVC was specifically driven by OVX+Veh mice. OVX+E2 abolished this relationship, restoring the size-independent dynamics characteristic of the estrogen-replete state. This pattern was reinforced by our multiple regression analyses that showed basal diameter significantly interacted with E2 to predict the magnitude of NVC enhancement (Table 2). Basal diameter therefore behaves not as a fixed property of the response but as a moderator with influence dictated by ovarian hormone status (or vice versa).

That the relationship between basal diameter and magnitude of NVC is moderated by ovarian hormone status rather than closely tracking vessel size indicates it is a physiological rather than a mathematically imposed effect. One possibility is that smaller PAs operate at higher maximal tone and retain greater capacity for NVC. A second possibility is that because NO is scavenged by hemoglobin more rapidly than by tissue, vessels with larger resting diameter degrade proportionally more NO, yielding smaller increases in smooth-muscle NO and less pronounced vasodilation^51^. A third possibility is that endothelial and neuronal machinery driving NVC, including endothelial caveolae^43^ and nNOS-expressing interneuron innervation^52^, are distributed unevenly along the arteriolar tree, as NVC dynamics are known to vary with depth and branch order within the vascular arbor^53^. Whichever mechanism predominates, estrogen decouples the magnitude of the hyperemic response from basal diameter. By extension, OVX would be expected to disadvantage areas perfused by larger PAs during heightened neuronal demand, a vulnerability E2 appears to correct. Our findings establish basal diameter as a moderator of NVC and underscore the importance of accounting for vessel size when comparing functional hyperemia across conditions that alter vascular morphology or hormonal state.

While the study has several important strengths, an *in vivo* approach to assess NVC in awake mice, a well-characterized model of metabolic dysfunction, and a surgical menopause and estradiol treatment, there are several limitations to consider. First, NVC examinations were performed using a cross-sectional approach. Tracking the same PAs before and after OVX and hormone treatment would allow more precise characterization of within-vessel dynamics over time. Second, our imaging approach assessed NVC exclusively through changes in PA diameter which, without concurrent measurement of neuronal activity, cannot distinguish the unique effects of E2 on the vasculature from its effects on neuronal or astrocytic activity upstream of the vascular response. Finally, as vasodilation of blood vessels in the S1BF are known to tightly track increases in local neuronal activity^54,55^, our use of whisker stimulation to measure changes in sensory-evoked PA diameter was chosen to interrogate NVC in PAs across treatment groups. After review of our data, one might suggest a longer term HFD model, or a Western high fat and sugar diet, might have resulted in more pronounced effects of metabolic disease on NVC. Countering this, other studies that also evaluated the effects of a diabetic phenotype in the S1BF agree that the homeostatic NVC response remains preserved despite hyperglycemia^33,54^, including one study using the Akita mouse model of chronic type 1 diabetes^39^. This provides strong evidence that if there is an effect of HFD-induced metabolic disease on PA NVC, it may manifest in other brain regions not examined here, such as the retina which may have greater vulnerability to hyperglycemia-induced vascular damage^56,57^ That said, neurovascular coupling is remarkably variable between brain regions, with some regions showing inverted or flat neuronal activity-vasodilation curves^58^. More mechanistic studies are needed to elucidate where in the brain NVC actually reflects metabolic state in order to develop better models to investigate the effects of hyperglycemia on cerebrovascular function.

Taken together, our findings advance the understanding of how menopause shapes cortical vascular dynamics. Additionally, they challenge the premise that pre-existing metabolic dysfunction occludes the neurovascular benefits of estrogen treatment and support the consideration of broader use of menopausal hormone therapy in mitigating risk of age-related cognitive decline.

## Methods

### Experimental timeline

Our study evaluated the effects of pre-existing metabolic dysfunction and subsequent E2 treatment on the NVC response to whisker stimulation in ovariectomized awake mice (Figure 1A). Mice underwent dual energy X-ray absorptiometry scans and glucose tolerance tests within one week prior to diet assignments to gather baseline readings at around 7.5 months of age (Pre-HFD), after 11 weeks of either HFD or CD to determine the consequences of HFD on metabolic health (Pre-OVX), and approximately 10 days after OVX to evaluate the interactions of diet and hormone treatment on metabolic health (Post-OVX). All mice were ovariectomized and treated with 17β-estradiol (E2) or vehicle (Veh) starting at approximately 10.5 months of age.

Chronic cranial window implantation was completed approximately three weeks prior to OVX or sham surgeries in all mice subjected to *in vivo* imaging. IOS imaging was performed within one week prior to OVX or sham surgeries in a subset of mice. NVC imaging was performed within one week prior to OVX or sham surgeries in one subset of mice (Pre-OVX/Sham) and approximately 4-6 weeks after OVX or sham surgeries in another subset of mice (Post-OVX/Sham). Upon completion of the experiments, at approximately 12 months of age, animals were deeply anesthetized with 5% isoflurane in an induction chamber. Deep anesthesia was confirmed by the absence of a response to firm toe pinch, after which the animals were immediately euthanized by decapitation using a rodent guillotine in accordance with approved standard operating procedures. Heart, kidneys, and uterine horn weights were collected. Uterine horn weights were collected to verify ovarian hormone depletion in ovariectomized mice receiving vehicle determined by uterine horn atrophy, and to verify bioavailability of estrogen in mice treated with E2 as given by uterine horn hypertrophy.

### Animals

Female wild type (C57BL6/J) and Thy1-GFP-M^28^ (bred on a C57BL6/J background) mice were used for all experiments. Mice were kept on a 12-hour light on/off cycle and given food and water *ad libitum*. All procedures were approved by the Tulane University Institutional Animal Care and Use Committee (protocol number 2295) and completed in accordance with the NIH Office of Laboratory Animal Welfare’s Public Health Service Policy on Humane Care and Use of Laboratory Animals and Guide for the Care and Use of Laboratory Animals. All procedures were performed in compliance with the ARRIVE guidelines.

### High-fat diet and control diet

At 10 weeks-old, all mice were switched from standard chow to a calorie-matched, phytoestrogen-free low fat control diet (CD; BioServ Control Diet F4031, Flemington, NJ; 16% kcal from fat). At seven and a half months of age, mice were randomly assigned to continue receiving CD or switched to an isoenergetic, phytoestrogen-free high fat diet (HFD; BioServ High Fat Diet F3282, Flemington, NJ; 59% kcal from fat). This HFD formulation was chosen because it has been shown to induce increased adiposity and glucose intolerance relative to CD^29–31^.

### Dual energy X-ray absorptiometry (DEXA) scans

Body composition scans were completed using dual energy X-ray absorptiometry (InAlyzer2, model S, Micro Photonics Inc.). The equipment was calibrated with a reference phantom as described by the manufacturer prior to use with experimental animals on a given day of scanning. Mice were anesthetized with isoflurane (5.0% for induction; 1.4-1.7% for maintenance). Fat mass (g) and lean mass (g) measurements were taken, and tibia lengths (mm) were recorded for normalizing uterine horn weights.

### Glucose tolerance tests (GTT)

Mice were removed from their home cages and separated into individual cages that included *ad libitum* access to water but without access to food for 4 hours. Blood samples were collected from the tail vein before an intraperitoneal injection of 20% glucose (2 g/kg body weight), and then again at 15, 30, 60, and 120 minutes following the glucose injection. Blood samples collected prior to glucose injection were used to determine fasting glucose levels, and together with the other timepoints to calculate area under the curve (AUC). Blood glucose levels were measured using a commercially available glucometer (OneTouch Verio Flex).

### Ovariectomy and hormone treatments

Mice were anesthetized with isoflurane (5.0% for induction; 1.4-1.7% for maintenance) and given a subcutaneous injection of meloxicam (5 mg/kg) prior to surgery. Bilateral ovariectomy surgery included incisions through skin and muscle to provide access to the intraperitoneal cavity, where ovaries were ligated and removed. Following this first surgical procedure, and while still under anesthesia, mice were implanted with a subcutaneous 5 mm silastic capsule (0.058 inch inner diameter and 0.077 inch outer diameter; Dow Corning,) on the dorsal aspect of the neck, which contained either 100% cholesterol vehicle (Sigma-Aldrich) or a mixture of 2.5% E2 (Sigma-Aldrich) with cholesterol vehicle, prepared in-house. Capsules were activated by submerging them in a sterile 0.9% NaCl solution inside a sterile tube to incubate at 37°C approximately 24 hours prior to surgical implantation. Capsules of these dimensions and estradiol concentration maintain blood plasma estradiol levels in the high physiological range^30^, and no replacements were required for the duration of our study as capsules have been demonstrated to remain active for up to five months post-implantation^32^. Each animal was allowed to reach full recovery for 1.5 weeks before completing subsequent procedures.

### Cranial window surgery

Cranial window surgery was performed to obtain optical access to the brain, as previously described, with a few minor changes. Mice were anesthetized with isoflurane (5.0% for induction; 1.4-1.7% for maintenance). In order to minimize inflammation and edema, mice were given subcutaneous injections of dexamethasone (0.2 mg/kg; VetOne) and carprofen (5.0 mg/kg, Zoetis, Inc) immediately prior to surgery. After placing mice on a stereotaxic frame (Stoelting), exposing the bone, and removing the overlying periosteum, two 00-96 x 1/16” screws (Protech Int.) were implanted into the skull of the contralateral hemisphere (frontal and parietal bone) to provide structural support for the permanent dental acrylic “headcap” used to affix mice to the *in vivo* rig during awake imaging. Using a pneumatic dental drill (Midwest Tradition) and with careful attention to not damage the dura mater, a 4 mm circular craniotomy was performed over the primary somatosensory cortex barrel field (S1BF; AP: −1.95, ML: +3.0). A 5 mm coverslip (#1; Electron Microscopy Sciences) was placed over the exposed brain and fixed to the skull with ethyl-cyanoacrylate glue (Krazy Glue). Lastly, a custom-designed two-pronged head bar was placed over the cranial window and secured to the skull with fast-curing dental acrylic (Lang Dental Mfg. Co., Inc). Each animal was allowed to reach full recovery for 2 weeks before completing subsequent procedures.

### Intrinsic optical signal imaging

We performed intrinsic optical signal (IOS) imaging to delineate the boundaries of the S1BF region within the cranial window, as previously described^33–35^. Briefly, mice were anesthetized (5% isoflurane for induction; <1% for maintenance) and head-fixed to a custom-built IOS rig using the implanted head bar. After capturing a reference image of the brain surface through the cranial window under green light (535 nm), contralateral whiskers were gently pressed onto a piezo bender actuator (Physik Instrumente) and affixed using dental wax. Intrinsic optical signals were captured under red light (630 nm) with a fast camera (Pantera 1M60; Dalsa) and frame grabber (64 Xcelera-CL PX4; Dalsa) using a custom-written MATLAB (MathWorks) routine. The imaging session consisted of 30 trials of whisker stimulation for 1.5 seconds in the rostro-caudal direction at 10 Hz with 20 second breaks, with images recorded at a depth of 300 ± 50 µm below the pial surface. Individual response signals were normalized to their respective baseline signal and summed to create a map of whisker stimulation-induced activity. This was then superimposed atop the vasculature reference image to identify the edges of the S1BF (Figure 1B, top). Once the functional map of this cortical area was repeatedly identified reliably in a subset of animals, the region was generalized to the rest of the experimental animals, and the position of cranial window placement remained consistent across animals.

### Habituation

Mice were habituated to the two-photon laser scanning microscopy (2PLSM) rig for four days before both the Pre-OVX and Post-OVX imaging sessions, as our paradigm to assess neurovascular coupling is performed in awake mice to avoid possible effects of anesthesia on vasodilation^36^. Mice were lightly anesthetized (4% isoflurane for 30 seconds) and secured to a custom-designed imaging platform using the implanted head bar. While head-fixed, each animal rested inside a clean metallic cylinder, and were allowed to fully wake up on the *in vivo* rig. Habituation consisted of individual periods of restraint while awake in total darkness across four consecutive days, at the following duration per day: 30 min, 60 min, 30 min, and 30 min. On days 3 and 4, the time spent in restraint was combined with 30 minutes of whisker stimulation.

### Two-photon laser scanning microscopy of neurovascular coupling

Experiments for the acquisition of NVC measurements were completed as previously described^33^ with minor modifications. 2PLSM imaging was performed using a custom-built microscope equipped with a Ti:sapphire laser (Chameleon Ultra II; Coherent) tuned to 910 nm, a 16x 0.8 NA water-immersion and long working distance objective (Nikon), and ScanImage software written in MATLAB. Mice were lightly anesthetized (4% isoflurane for 30 seconds), and given a retroorbital 100 µL injection of fluorescein isothiocyanate-dextran 70 kDa (FITC, 5% w/v in sterile saline) to label blood serum, which was used to visualize the cerebral vasculature. Immediately after injection, mice were secured to the *in vivo* rig and positioned beneath the microscope objective (Figure 1B, bottom). Mice were allowed to fully recover from anesthesia for 15-30 minutes before the start of the imaging session. Penetrating arterioles (PAs) were selected based on whether they fell within the boundaries of the S1BF as given by IOS+vasculature maps, dilated in response to air puffs, and agnostic to basal diameter and hormone treatment. At a depth of 50 ± 20 µm from the pial surface, cross sections of identified PAs were imaged at 9.45 Hz for a total duration of 25 seconds per trial (Figure 1C, left). For each trial, air puffs generated with a picospritzer III (4Hz, 5 s, 20 psi, 30 s/cycle) were used to stimulate the whiskers contralateral to the imaged S1BF, which were initiated 10 seconds after onset of image acquisition. An Arduino-based custom-built I/O controller was used to synchronize imaging acquisition to the onset of whisker stimulation. Two to six PAs were identified and imaged per mouse, and ten stimulation trials were performed one minute apart for each PA and averaged to minimize variability due to artifacts inherent to *in vivo* awake mouse imaging.

### Image analysis of NVC responses

For each stimulation trial, PA dilation dynamics were quantified using a custom-written routine in MATLAB^33^. Briefly, the image intensity of each PA cross-section was adjusted, binarized, and as the resulting mask takes the form of an ellipsis, the minor axis (i.e. the minimum diameter running through the centroid of the PA) was calculated for every frame recorded and will henceforth be referred to as the “diameter” of the PA at a given frame (Figure 1C, right). Basal diameter was calculated as the average diameter from second 3 to the onset of the whisker stimulation at second 10, which was done to avoid anticipatory vasodilation artifacts frequently observed within the first 1-3 seconds of each trial^33^. The traces from the 10 trials per PA were averaged to reduce variability from artifacts related to awake imaging. NVC diameter response was evaluated between seconds 10 and 15, corresponding to the whisker stimulation period, and the basal diameter calculation was used to normalize stimulation-induced changes in diameter per frame across a given trial (basal diameter being considered 100%). Several parameters of NVC were then obtained from these recorded traces (Figure 1D): the averaged PA diameter from seconds 3 to 10 before whisker stimulation onset, in µm (basal diameter); the maximum % change in PA diameter during whisker stimulation relative to basal diameter (amplitude); the time in seconds after whisker stimulation onset to reach maximum amplitude (t_Max_); the area under the curve calculated during whisker stimulation (AUC); the maximum diameter of each vessel subtracted by its minimum diameter across the time series in µm (dynamic range).

### Statistical analysis

Animals were randomly assigned to diet and hormone treatments, and experimenters were blinded to hormone treatment for the duration of each experiment. Statistical analyses were performed and figures plotted using GraphPad Prism Version 11.0.2. Data from mice with uterine horn weights that were incongruent with their randomized hormone assignments were excluded from all analyses. Results from DEXA and GTTs were analyzed cross-sectionally at each timepoint they were completed (i.e. Pre-HFD, Pre-OVX, and Post-OVX) using two-tailed Student’s *t* tests or ordinary two-way ANOVAs followed by *post hoc Šídák tests* (which correct for multiple comparisons). For the analysis of NVC responses, recordings were taken from two to six PAs per mouse. NVC parameters were analyzed cross-sectionally using the two most responsive PAs from each mouse, as given by the PAs with the highest whisker stimulation-evoked amplitude values. This procedure minimized sampling bias and potentially uneven contribution of mice with different numbers of imaged PAs. Average t_Max_ and standard deviations (SD) were calculated across treatment groups at a given experimental NVC imaging timepoint (i.e. Pre-OVX/Sham or Post-OVX/Sham), and PAs with a t_Max_ greater than or equal to 2 SD from the timepoint average were excluded from all analyses. NVC parameters were analyzed per PA using two-tailed Student’s *t* tests, ordinary two-way ANOVAs followed by *post hoc Šídák* tests. Multiple linear regression analyses were used to evaluate main and interaction effects while controlling for the effects of basal diameter, and *R*, *R*^2^, adjusted *R*^2^, and results from each model’s omnibus *F* test were reported, in addition to *t* for each covariate. In these instances, absolute changes in NVC amplitude were used (instead of % change from basal diameter) to minimize parameter covariance. *Cohen’s d*, partial *eta* squared (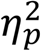), and unstandardized regression coefficients (*B*) are reported alongside all results of significance testing to estimate the magnitude of observed effects^37^. *Cohen’s d* was calculated using a publicly available effect size calculator (https://lbecker.uccs.edu), partial *eta* squared was computed manually, and *B* was included as part of GraphPad Prism’s regression analyses results. Data are presented as mean ± SD unless otherwise indicated in figure captions, with significance determined at *p*<0.05. In the results and figures, level of significance is denoted by the following conventions: \**p*<0.05; \*\**p<*0.01; \*\*\**p*<0.001; \*\*\*\**p*<0.0001

## Acknowledgements

The authors thank our Program Project Grant co-investigators Dr. Jill Daniel, Dr. Sarah Lindsey, Dr. Laura Schrader, and Dr. Andrea Zsombok for their guidance in the development and coordination of resources to support this project.

## Author contributions

ZMTP and RM conceived the study. ZMTP, JCI, IFU, ABW, IP, and HT performed the experiments. ZMTP, JCI, JBR, and RM analyzed the data. ZMTP and RM wrote the paper. RM oversaw the experiments. All authors read and approved the final version of the manuscript.

## Material availability

The datasets generated during and/or analyzed during the current study are available from the corresponding author upon reasonable request. MATLAB codes for neurovascular coupling analysis, and custom-design pieces are available at https://github.com/mostanylab/NVC.

