## Supplementary Figure 1 for "Estradiol enhances sensory-evoked dilation of cortical penetrating arterioles in a mouse model of menopause despite pre-existing metabolic dysfunction"

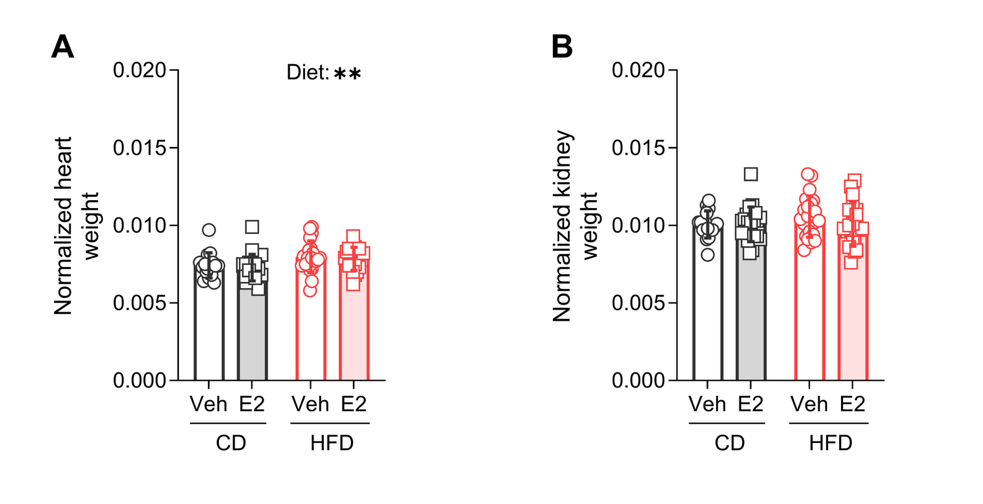

**Supplementary Figure 1.**

Comparison of (A) normalized heart weight from CD/OVX Veh-treated mice (n=16 mice), CD/OVX E2-treated mice (n =21 mice), HFD/OVX Veh-treated mice (n=21 mice), and HFD/OVX E2-treated mice (n=20 mice); (B) normalized kidney weight from CD/OVX Veh-treated mice (n=16 mice), CD/OVX E2-treated mice (n =21 mice), HFD/OVX Veh-treated mice (n=22 mice), and HFD/OVX E2-treated mice (n=20 mice).

**p<0.01. Veh: vehicle; E2: estradiol; CD: control diet; HFD: high-fat diet
